# Response of soil microbial communities to alpine meadow degradation severity levels in the Qinghai-Tibet Plateau

**DOI:** 10.1101/490375

**Authors:** Wenjuan Zhang, Xian Xue, Fei Peng, Quangang You, Jing Pan, Chengyang Li, Chimin Lai

## Abstract

Soil microbial community structure is an effective indicator to reflect changes in soil quality. Little is known about the effect of alpine meadow degradation on the soil bacterial and fungal community. In this study, we used the Illumina MiSeq sequencing method to analyze the microbial community structure of alpine meadow soil in five different degradation levels (i.e., non-degraded (ND), slightly degraded (LD), moderately degraded (MD), severely degraded (SD), and very severely degraded (VD)) in the Qinghai-Tibet Plateau. *Proteobacteria, Actinobacteria*, and *Acidobacteria* were the mainly bacterial phyla in meadow soil across all five degradation levels investigated. *Basidiomycota* was the mainly fungal phylum in ND; however, we found a shift from *Basidiomycota* to *Ascomycota* with an increase (severity) in degradation level. The overall proportion of *Cortinariaceae* exhibited high fungal variability, and reads were highest in ND (62.80%). Heatmaps of bacterial genera and fungal families showed a two-cluster sample division on a genus/family level: (1) an ND and LD group and (2) an SD, VD, and MD group. Redundancy analysis (RDA) showed that 79.7%and 71.3% of the variance in bacterial and fungal composition, respectively, could be explained by soil nutrient conditions (soil organic carbon, total nitrogen, and moisture) and plant properties (below-ground biomass). Our results indicate that meadow degradation affects both plant and soil properties and consequently drives changes in soil microbial community structure.

## Introduction

The Qinghai-Tibet Plateau (QTP) has an area over 250×10^4^ km^2^ within China and is the highest plateau in the world with an average elevation of ~4500 m a.s.l. [1]. Alpine meadows comprise approximately 38% of all grassland area in the QTP and are the primary ecosystem utilized by the Tibetan people and their livestock [2,3]. Accordingly, the alpine meadow is considered to be one of the most critically important ecosystems in the QTP. The ecological functions of alpine meadow ecosystems in the QTP are also important, which include water storage [4], biodiversity maintenance [5], and soil carbon (C) sequestration [6].

Furthermore, the lower air temperature and higher altitude make alpine meadows more sensitive to global warming. Thus, these ecosystems are considered as good indicators of environmental change [2]. In recent decades, frequent reports on the degradation of alpine meadows in the source regions of the Yangtze and Yellow rivers have been attributed to climate warming and anthropogenic activities [7,8]. The alpine grassland degradation has led to a variety of ecological consequences, including alterations plant community composition, decreased plant species richness and biomass [9], and accelerated soil erosion [10].

Soil bacteria and fungi play crucial roles in soil nutrient supplies and element cycling in terrestrial ecosystems [11], and their composition and diversity are sensitive to disturbances [12,13]. Different microbes exhibit various ability to efficiently utilize soil organic matter (SOM) and the composition of microbial decomposers directly influence a variety of ecosystem processes, such as CO2 flux and litter decomposition. Previous studies have shown that soil microbial communities are affected by plant characteristics and soil properties [14,15]. For example, vegetation type has a strong effect on soil microbial communities in determining the physical soil environment and the availability of nutrients [16]. Soil substrate availability and heterogeneity are important factors responsible for changes in microbial communities [17,18]. Potential changes in soil nutrient availability [19], soil moisture [20], and plant composition and biomass during processes of grassland degradation [21] would inevitably alter the composition and diversity of soil microbial communities. The current literature on alpine meadow degradation mainly focuses on plant and soil characteristics; however, knowledge regarding the effects of meadow degradation on soil microbial communities and their diversity remains largely inadequate [22]. Therefore, it is essential to understand how the composition and diversity of microbial communities respond to alpine meadow degradation in the QTP and to consider the key influencing factors and to provide important insights for the alpine meadow health assessment and management.

In this study, we selected alpine meadows under different levels of degradation severity (i.e., non-degraded (ND), slightly degraded (LD), moderately degraded (MD), severely degraded (SD), and very severely degraded (VD)), applying the space-for-time substitution method [9]. We used the Illumina MiSeq sequencing method (Illumina, Inc., USA) to determine soil bacterial and fungal composition and their diversity with changes in edaphic and plant properties under degradation. The specific aims of this study were: 1) to investigate how soil bacterial and fungal communities vary with changes in explicit soil and plant properties in the QTP, and 2) to discern which factors under conditions of alpine meadow degradation significantly influence associative microbial community structure.

## Materials and methods

### Study area

The study area was located at the source of the Yangtze River on the QTP (34°49’N, 92°55’; 4635 m a.s.l.) (Fig 1). The annual average mean air temperatures are -3.8°C. Mean annual precipitation is 285 mm with greater than 93% falling during the warm growing season (April–October) [2]. The site is used for grazing during the summer. The typical vegetation is alpine meadow dominated by *Kobresia capillifolia, Kobresia pygmaea*, and *Carex moorcroftii* [3].

**Fig 1.**
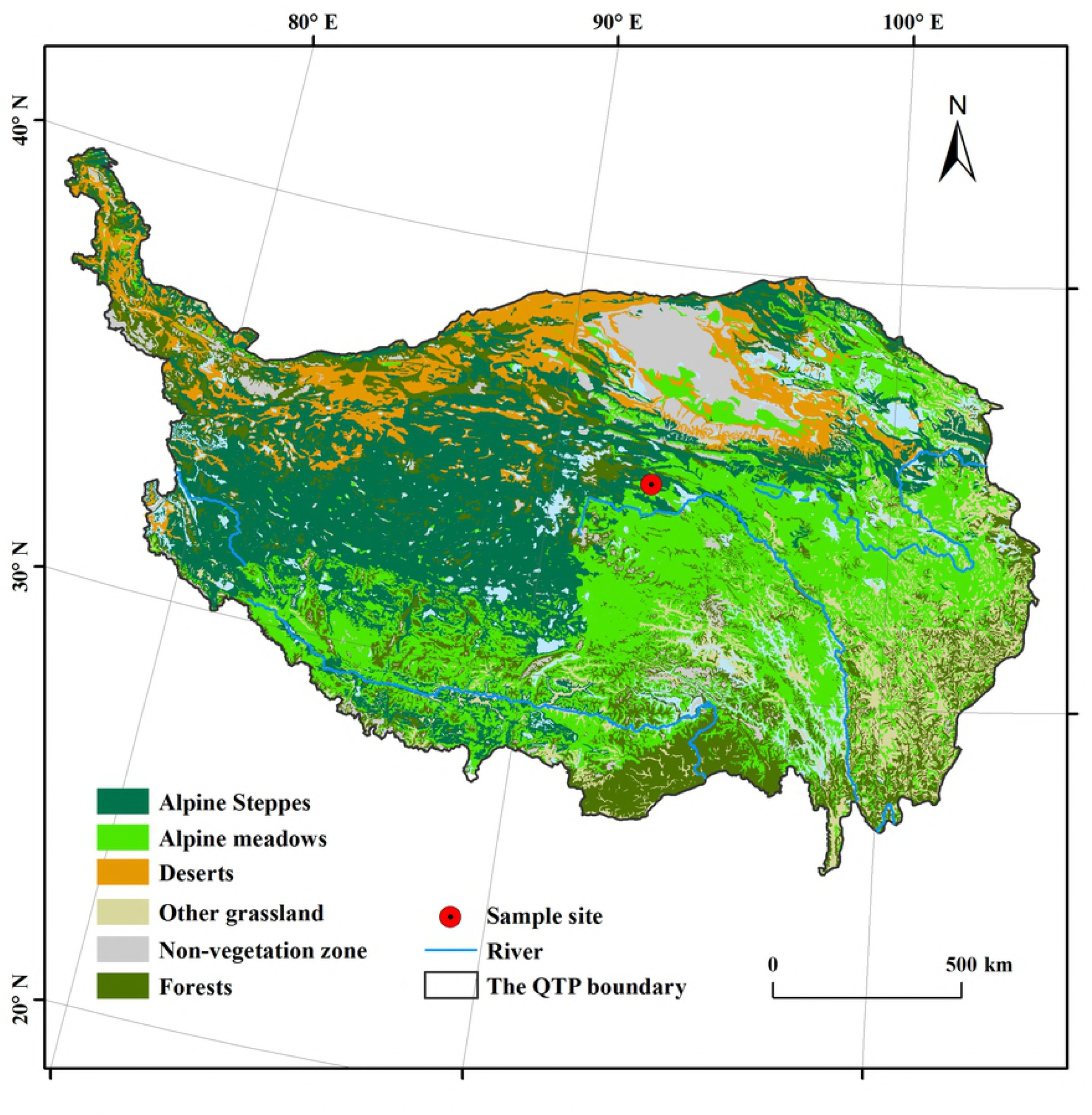
The location of the sampling site.

The ND, LD, MD, SD, and VD alpine meadow sites were chosen based on vegetation coverage as described by Liu et al. (2018) [23]. Vegetation coverage percentages are as follows: 80%~90% for ND, 70%~80% for LD, 50%~70% for MD, 30%~50% for SD, and <30% for VD.

### Plant measurement and soil sampling procedures

Five quadrats (5 m × 5 m) at a spatial distance of greater than 50 m were randomly selected at each of the five alpine meadow sites. Three plots were sampled from each quadrat. Plant cover, height, and composition of each quadrat were measured using point-intercept sampling, employing a 30 cm × 30 cm square frame, with 100 sampling points spaced equidistantly within the frame [24]. Plant height was estimated by randomly measuring 10 individuals in each quadrat. Following this, plants were clipped at the soil surface, and harvested plants were categorized into different functional groups: graminoids, sedges, and forbs. Aboveground and belowground plant biomass (abbreviated as AGB and BGB in this study, respectively) were separated, dried in an oven at 65°C for 48 h before being weighed.

On July 3, 2017, three 50 mm diameter surface soil cores (from the top 10 cm soil layer) were randomly collected from each quadrat. These soil cores were then mixed, homogenized, and sieved (<2 mm) to remove roots and other plant material [25]. Each soil sample was placed in a sterile centrifuge tube before being immediately transported to the laboratory and stored at -80°C for total DNA extraction and molecular analyses.

From each quadrat, three soil profiles were randomly collected for soil sampling (0 to 10 cm). The soil samples were measured for soil bulk density (BD), soil moisture (SM), and soil nutrient content piling at each of its four corners.

### Soil characterization

Soil BD was measured using 100 cm^3^ rings and calculated from dry soil matter. At the same time, SM was measured by drying soil samples taken from the rings at 105°C for 48 h. Soil organic carbon (SOC) was determined using the potassium dichromate oxidation titration method [26]. Total nitrogen (TN) in soil was determined using the semimicro Kjeldahl digestion procedure [2]. Ammonium-nitrogen (NH_4_^+^-N) and nitrate nitrogen (NO_3_^-^-N) content in soil were measured using the UV-3300 Spectrophotometer (Mapada Instruments Co., Ltd., Shanghai, China).

### DNA extraction, Polymerase chain reaction and Illumina sequencing

DNA was extracted from 0.5 g soil samples using the PowerSoil DNA Isolation Kit (Mo Bio Laboratories, Carlsbad, CA, USA) according to the manufacturer’s instructions. The universal primer pair 338F/806R [27] for bacteria and ITS1 [28] for fungi were used for amplification and MiSeq sequencing of the polymerase chain reaction (PCR) products. PCR amplification was conducted in a 25 μL reaction system using TransGen AP221-02 (TransGen Biotech, Beijing, China) and carried out in an ABI GeneAmp 9700 (Applied Biosystems, Inc., Carlsbad, USA). After purified and quantitated, the PCR products were sequenced using the Illumina Miseq platform.

The 16s and ITS1 rRNA gene sequences associated this study were submitted to the National Center for Biotechnology Information (NCBI) Sequence Read Archive (SRA) (accession no. PRJNA490659).

### Data analysis

The QIIME (version 1.8.0) was used to estimate α-diversity (i.e. Chao1, Shannon, and Good’s coverage) for each sample [29]. The changes of species composition under different degradation stages, principal component analysis (PCA), and heatmaps were conducted using the R language (3.2.0). One-way analysis of variance (ANOVA) tests was used to evaluate significant levels for all factors analyzed herein. The SPSS 17.0 software package (SPSS Inc., Chicago, IL, USA) was used to calculations and analyses. The CANOCO software (version 4.5) was used to run redundancy analysis (RDA) [30].

## Results

### Response of plant and soil factors to alpine meadow degradation

Plant coverage in MD, SD, and VD decreased significantly (*P* < 0.05) by 32.48%, 17.52%, and 84.67% compared to ND and by 38.13%, 24.42%, and 85.95%, respectively, compared to LD (Table 1). Plant AGB in MD, SD, and VD significantly (*P* <0.05) decreased by 47.09%, 51.60%, and 84.14%, respectively, compared to LD (Table 1). Plant BGB in SD and VD also significantly decreased by 60.43% and 67.12% compared to ND and by 59.67% and 66.49% in SD and compared to LD, respectively (Table 1). The ND and LD sites not exhibited a significant differ in plant coverage, AGB, or BGB (Table 1). There was no significant change in species richness (SR) and plant height throughout the whole degradation process (Table 1). For plant group, sedge biomass in SD and VD significantly decreased compared to LD, but we found no significant change in the biomass/proportion of graminoids and forbs as degradation severity increased (Table 2).

**Table 1.** Properties of plant community at different degraded alpine meadow (n=3).

| Items | ND | LD | MD | SD | VD |
| --- | --- | --- | --- | --- | --- |
| **Plant coverage (%)** | 91.33±1.78ab | 99.67±0.27a | 61.67±5.04c | 75.33±6.15bc | 14±4.49d |
| **Plant height (cm)** | 1.63±0.27a | 2.47±0.08a | 1.66±0.06a | 2.32±0.33a | 1.89±0.17a |
| **Species richness (No./m<sup>2</sup>)** | 6.67±0.72a | 7±0.82a | 6±0.82a | 6±0.47a | 7±0.47a |
| **Above-ground biomass (g/m<sup>2</sup>)** | 241.63±1.81ab | 383.85±43.19a | 203.11±4.94bc | 185.78±50.39bc | 60.86±15.39c |
| **Blow-ground biomass (g/m<sup>2</sup>)** | 1918.45±48.36a | 1882.14±310.92ab | 861.30±205.47abc | 759.05±25.54bc | 630.79±237.42c |

Note: The data at the table are showed by average value ± standard error. Data with the different letter in same row are significantly different (P<0.05). The same below.

**Table 2.**
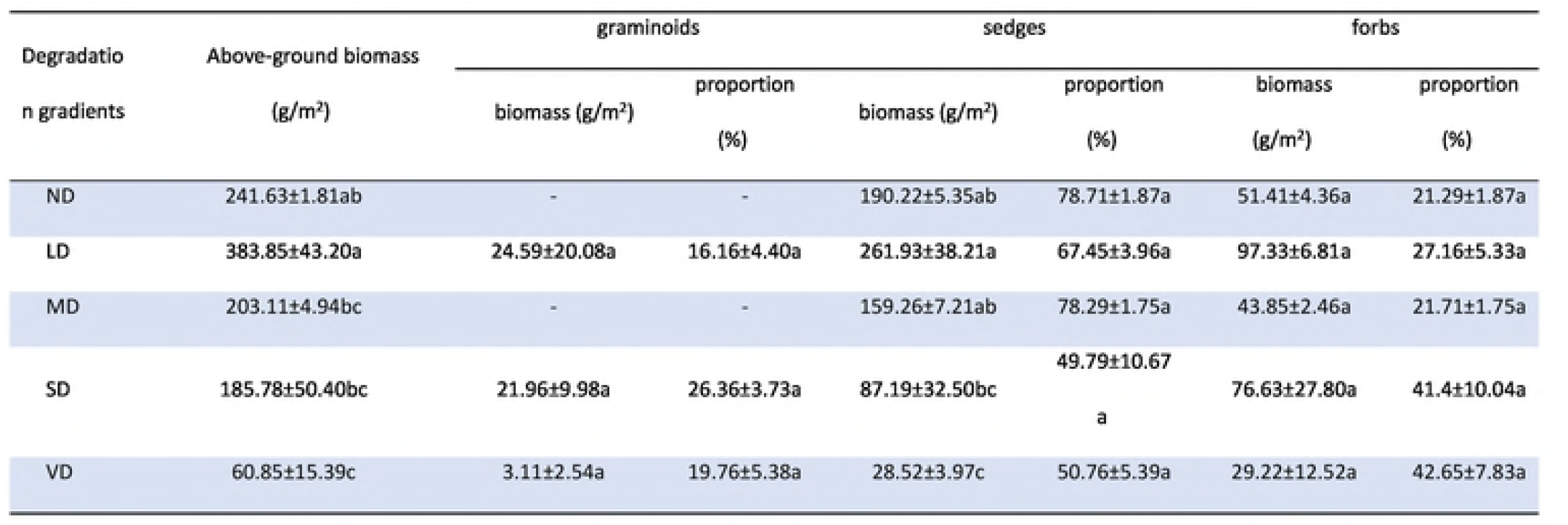
Properties of plant AGB and proportion at different degraded alpine meadow (n=3).

Soil TN, SOC, N0_3_^−^-N, NH_4_^+^-N, and SM content decreased significantly in MD, SD, and VD compared to ND, but BD exhibited the opposite response pattern (Table 3). In MD, there were significant decreases in soil TN, SOC, NO_3_^-^-N, NH_4_^+^-N, and SM by 62.20%, 67.75%, 48.40%, 47.94%, and 43.57%, respectively, compared to ND (Table 3); in SD, the content of soil TN, SOC, NO_3_^-^-N, NH_4_^+^-N, and SM decreased significantly by 65.35%, 67.11%, 60.34%, 50.09%, and 65.66%, respectively, compared to ND (Table 3); in VD, there were significant decreases in soil TN, SOC, NO_3_^-^-N, NH_4_^+^-N, and SM by 70.08%, 76.16%, 57.64%, 43.25%, and 66.72%, respectively, compared to ND (Table 3). However, soil BD increased significantly (*P* < 0.05) by 37.14%, 28.57%, and 47.14% in MD, SD, and VD, respectively, compared to ND (Table 3). We found no significant difference in SOC, TN, NO_3_^-^-N, NH_4_^+^-N, SM, and BD between ND and LD (Table 3).

**Table 3.** Properties of soil at different degraded alpine meadows (n=3, depth=0-10cm).

| Items | ND | LD | MD | SD | VD |
| --- | --- | --- | --- | --- | --- |
| Organic carbon (SOC, g/kg) | 15.69±1.10a | 17.95±0.42a | 5.06±0.62b | 5.16±0.29b | 3.74±0.49b |
| Total nitrogen (TN, g/kg) | 1.27±0.06a | 1.4±0.03a | 0.48±0.03b | 0.44±0.01b | 0.38±0.03b |
| Nitrate nitrogen (mg/kg) | 14.07±0.23b | 16.37±0.56a | 7.26±0.55c | 5.58±0.12c | 5.96±0.18c |
| Ammonium nitrogen (mg/kg) | 16.81±0.64a | 14.73±0.22a | 8.75±0.60b | 8.39±0.23b | 9.54±0.78b |
| Soil moisture content (%) | 27.61±0.69a | 23.46±0.42a | 15.58±0.84b | 9.48±0.79c | 9.19±1.00c |
| Soil bulk density (g/cm<sup>3</sup>) | 0.70±0.01b | 0.67±0.02b | 0.96±0.03a | 0.90±0.04a | 1.03±0.02a |

### α-diversity indices based on MiSeq sequencing

For bacteria, MD yielded the highest richness value (Chao1 = 1187.75), while SD yielded the lowest (Chao1 = 1086.23). The Shannon index not only provides simple SR (the number of species present) but also the level of abundance of each species (species evenness) as distributed among all species in a community. ND yielded the highest diversity value (Shannon = 8.46) among the five samples, and SD yielded the lowest diversity value (Shannon = 8). From 962 to 1104 OTUs were detected in total for all samples at a 3% genetic distance. The progression of rarefaction curves (99.01%−99.26%; Good’s coverage) was very close for all samples (S1 Table).

For fungi, LD also yielded the highest richness value (Chao1 = 645.85), followed by VD (Chao1 = 575.29), and ND yielded the lowest richness value (Chao1 = 517.94). VD yielded the highest diversity value (Shannon = 6.38), however, ND yielded the lowest diversity value (Shannon =3.13). The number of OTUs ranged from 423 to 564 in the samples, for which LD had the highest one and ND the lowest (S1 Table).

### Taxonomic composition based on MiSeq sequencing

Bacterial OTUs could be assigned to 28 phyla, 156 families, and 170 genera. In total, 20 different phyla (*Proteobacteria, Acidobacteria, Actinobacteria, Bacteroidetes, Chloroflexi, Gemmatimonadetes, Verrucomicrobia*, etc.) out of the 28 total bacterial phylotypes were common to the five libraries (Fig 2A), contributing from 99.84%, 99.62%, 99.72%, 99.91%, and 99.74% of the total reads in the ND, LD, MD, SD, and VD libraries, respectively. *Proteobacteria* was the most abundant division, comprising 26.48% (350) of the OTUs and 44.18% (10 078) of the reads across all samples. *Acidobacteria*, the second most abundant phylum, comprised 14.83% (196) of the OTUs and 18.44% (4213) of the reads in all libraries. These two phyla collectively accounted for 60.88%, 66.83%, 64.31%, 63.65%, and 57.42% of the total reads in the ND, LD, MD, SD, and VD libraries, respectively (Fig 2A). However, the proportion of *Proteobacteria* exhibited low variability in the different samples, namely, ND (46.63%; 10 224 reads), LD (46.80%, 11,442 reads), MD (39.08%, 8,069 reads), SD (49.68%, 10,637 reads) and VD (38.71%, 10,021reads). Additionally, reads from *Acidobacteria* fluctuated in the different samples, for which the proportion of reads was highest in MD (25.23%; 5210 reads).

**Fig 2.**
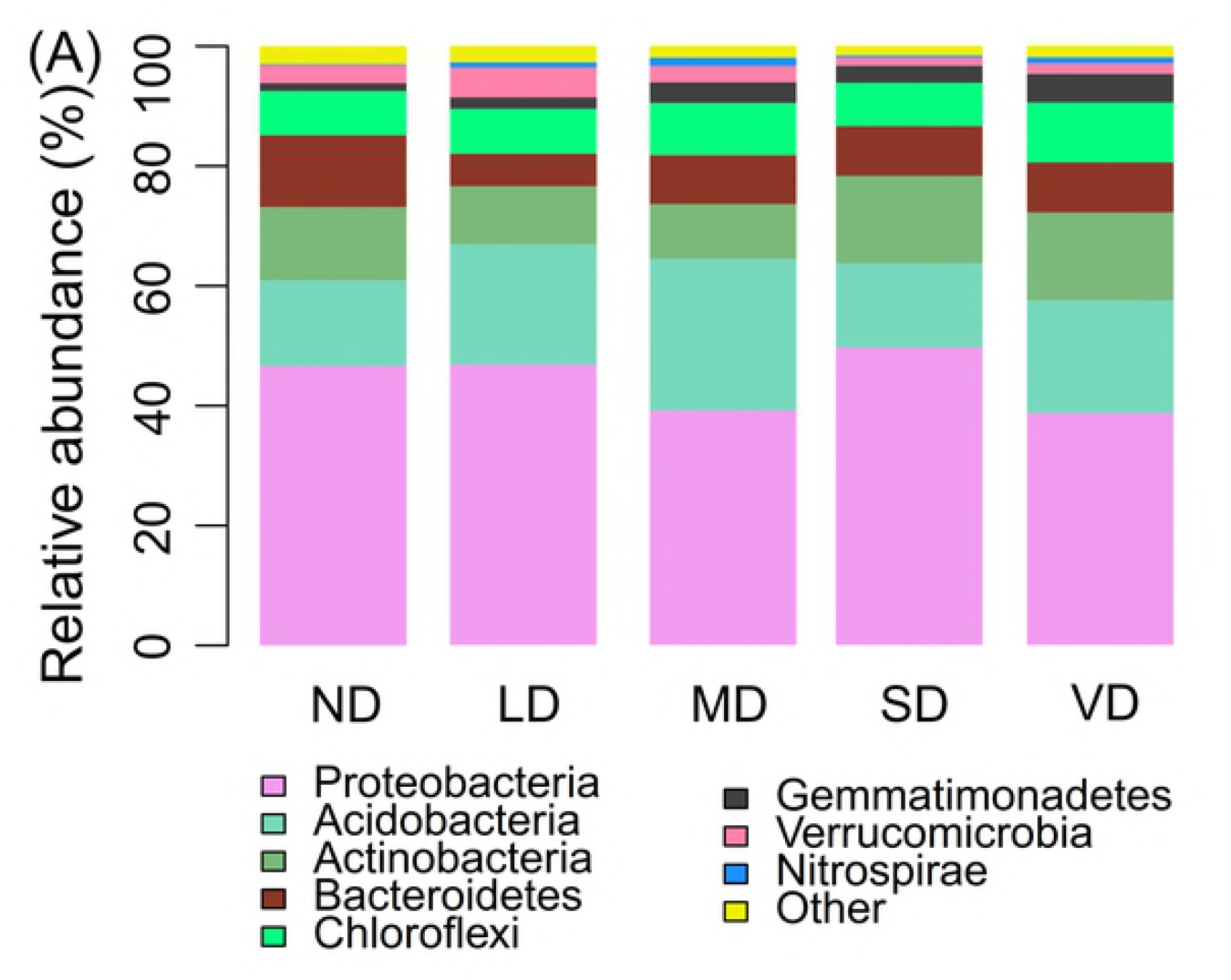
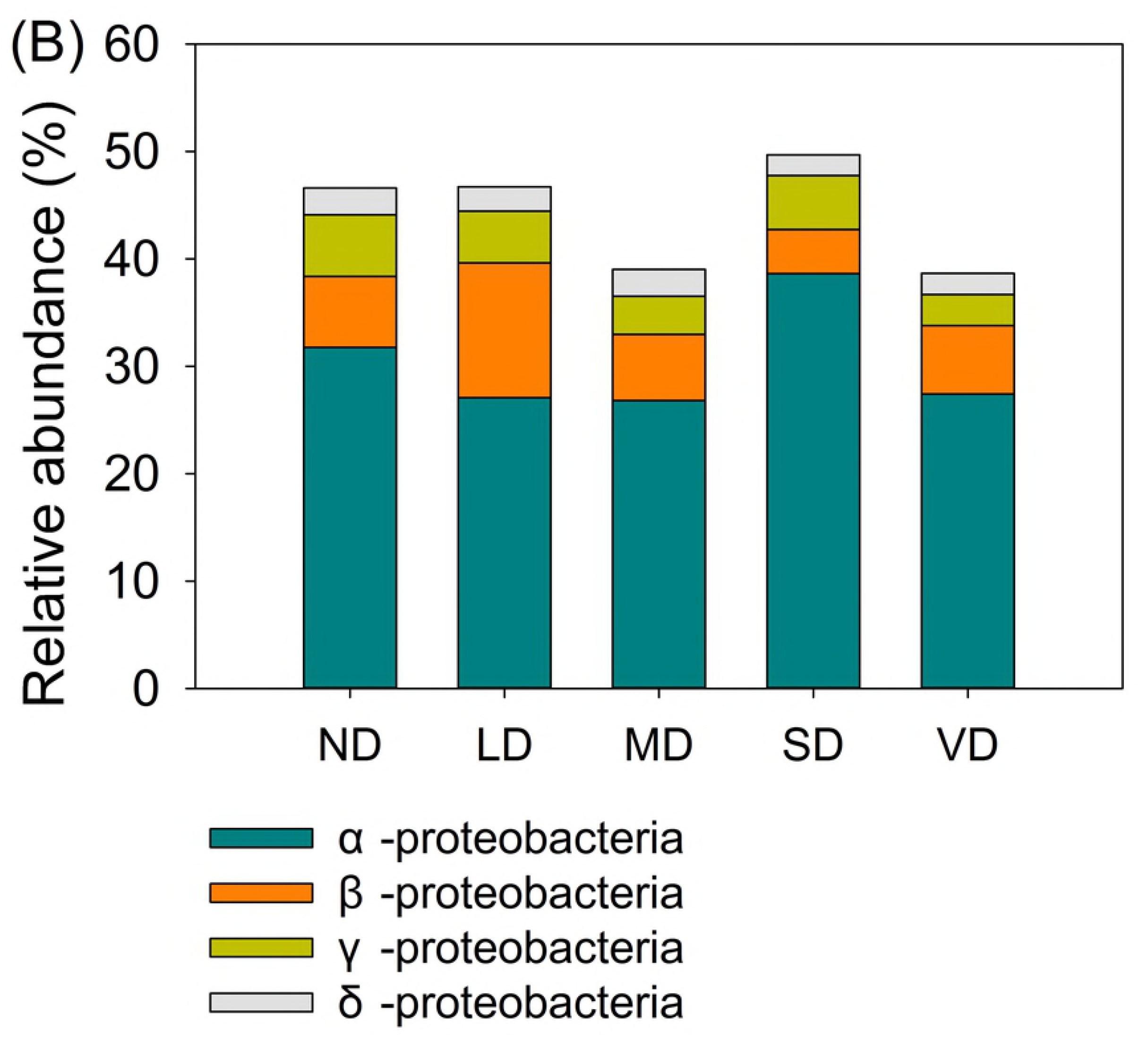
Relative abundance of the bacterial community at the phylum (A) level and class distribution of proteobacteria (B).

Although members of the *α-proteobacteria* class dominated the *proteobacteria* phylum (10.82% and 143 OTUs) in all libraries, they accounted for 30.34% of the total reads (Fig 2B). The subdivision of *β-proteobacteria, δ-proteobacteria*, and *γ-proteobacteria* classes comprised of 7.16%, 4.42%, and 2.21% of the total reads, respectively.

The fungal communities were assigned to 8 phyla, 78 families, and 138 genera. *Ascomycota* was the most dominant division, comprising 34.52% (339) of the OTUs and 44.43% of the total reads (Fig 3A). *Basidiomycota* was the second largest division, with 12.63% (124) of the OTUs and 29.92% of the total reads. These two phyla collectively accounted for 95.88%, 72.04%, 64.78%, 74.50%, and 64.55% of the total reads in ND, LD, MD, SD, and VD, respectively (Fig 3A). However, *Ascomycota* exhibited high variability in read abundance of the different samples; namely, in increasing order of abundance, ND (14.47%; 5641 reads), LD (37.90%; 13 649 reads), MD (48.06%; 20 251 reads), SD (63.35%; 24 980 reads), and VD (58.38%; 23 434 reads). In contrast, *Basidiomycota* reads decreased with an increase in degradation severity; accordingly, ND had the highest read value (81.40%; 31 729 reads). Reads from *Zygomycota* and *Glomeromycota* fluctuated in the different samples; namely, VD had the highest read value (Fig 3A).

**Fig 3.**
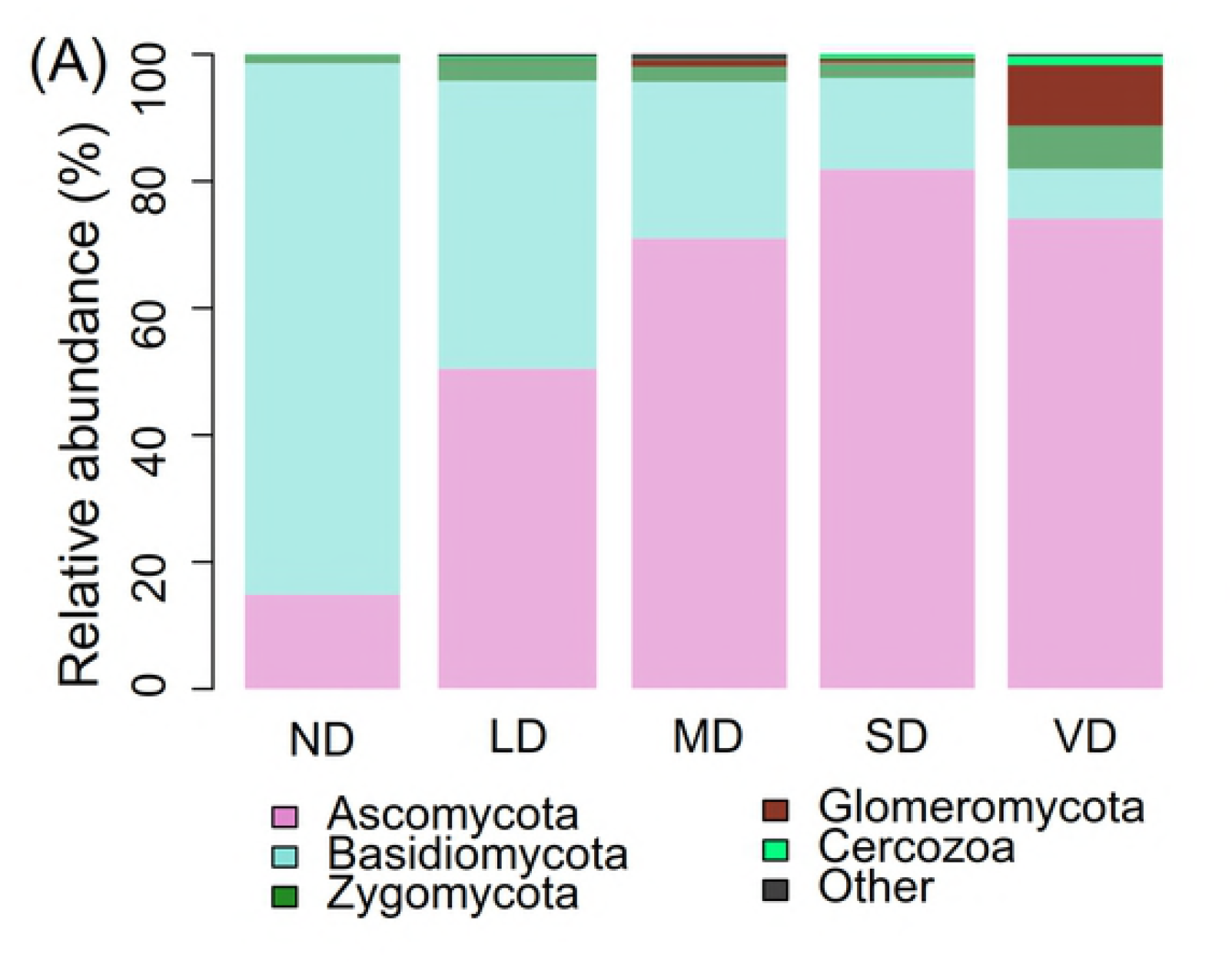
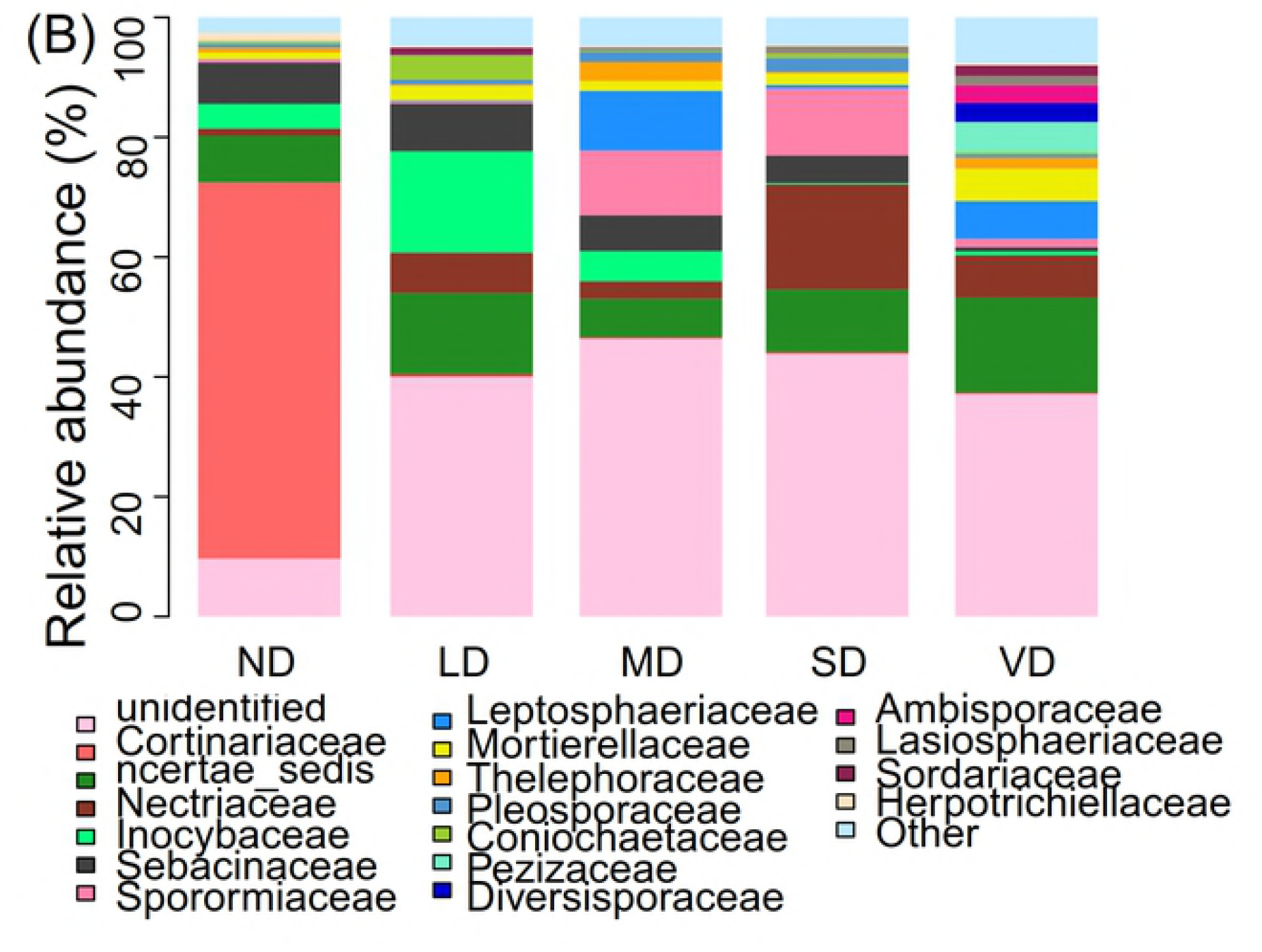
Relative abundance of the fungal community at the phylum (A) and family(B) level.

On a family level, we found abundant unidentified fungal sequences in each sample. The total relative abundance of unidentified fungi in ND, LD, MD, SD, and VD were 9.65%, 40.06%, 46.40%, 43.89%, and 37.19%, respectively (Fig 3B). *Cortinariaceae* members dominated the Basidiomycota phylum, and the proportion of reads exhibited high variability in read abundance of the different samples; namely, ND (62.80%; 24 476 reads), LD (0.42%; 150 reads), MD (0.23%; 98 reads), SD (0.26%; 101 reads), and VD (0.22%; 90 reads). *Incertae-sedis*, the third most abundant group (7.84%; 77 OTUs), comprised 10.78% (4208) of the reads in all libraries. These three groups combined, namely, unidentified, *Cortinariaceae*, and *incertae-sedis*, comprised of greater than 53.21% of the total sequences from all five sample sites.

### Taxonomic composition based on MiSeq sequencing

In order to analyze microbial community similarity among the five mixed samples, we generated heatmaps applying hierarchical cluster analysis. For bacteria, the heatmap (Fig 4A) was based on the top 20 abundant bacterial genera. The heatmap showed a two-sample cluster division. The first cluster was the ND and LD group, and the second cluster was the SD and VD group, which first clustered together before clustering with MD, resulting in the second SD, VD, and MD group. Results from PCA also showed that the bacterial communities of SD and VD grouped to the left of the graph along the PC1 axis, accounting for 41.08% of total variation, whereas ND and LD grouped along the PC2 axis, with a total variance of 31.92% (Fig 5A).

**Fig 4.**
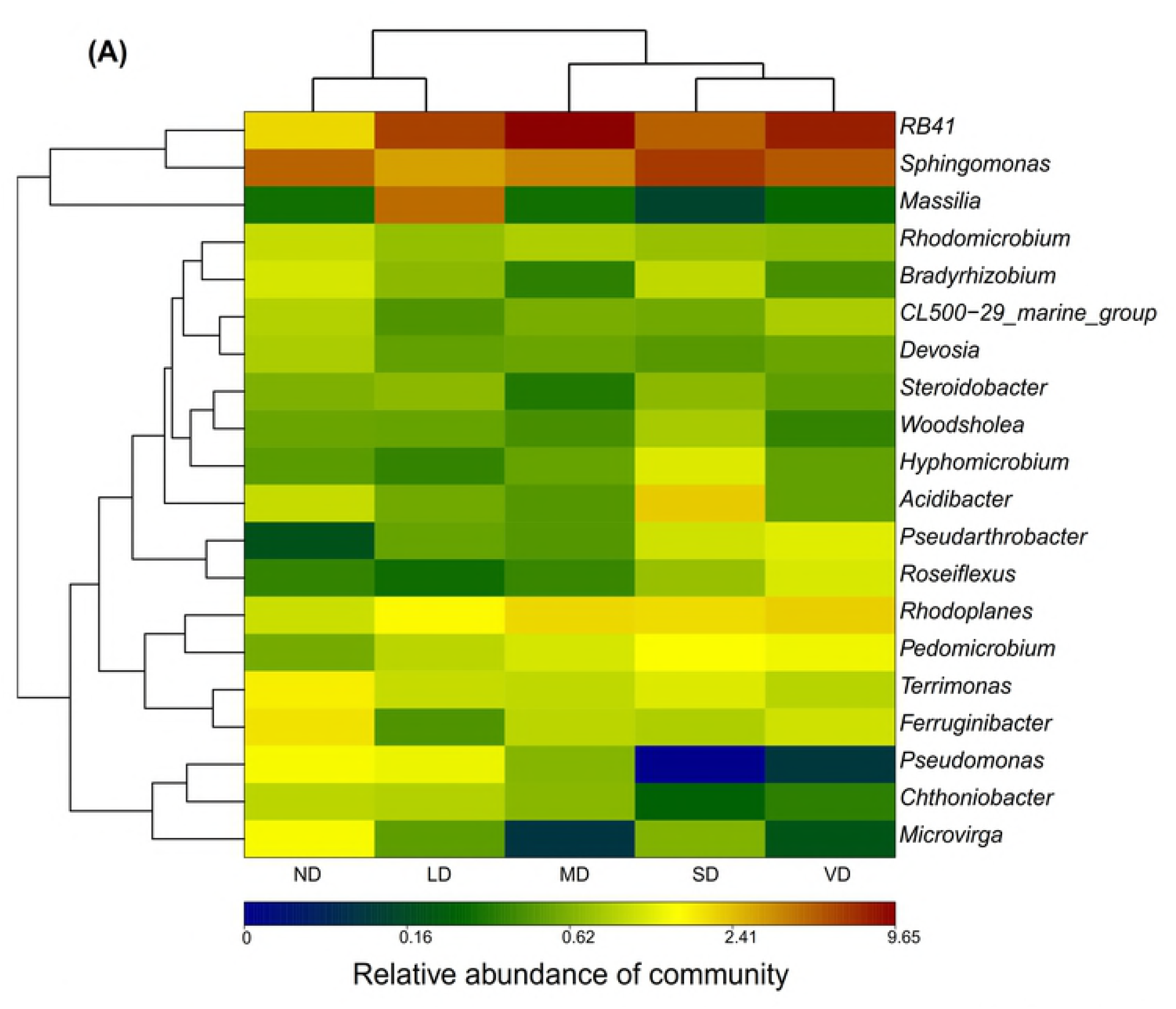
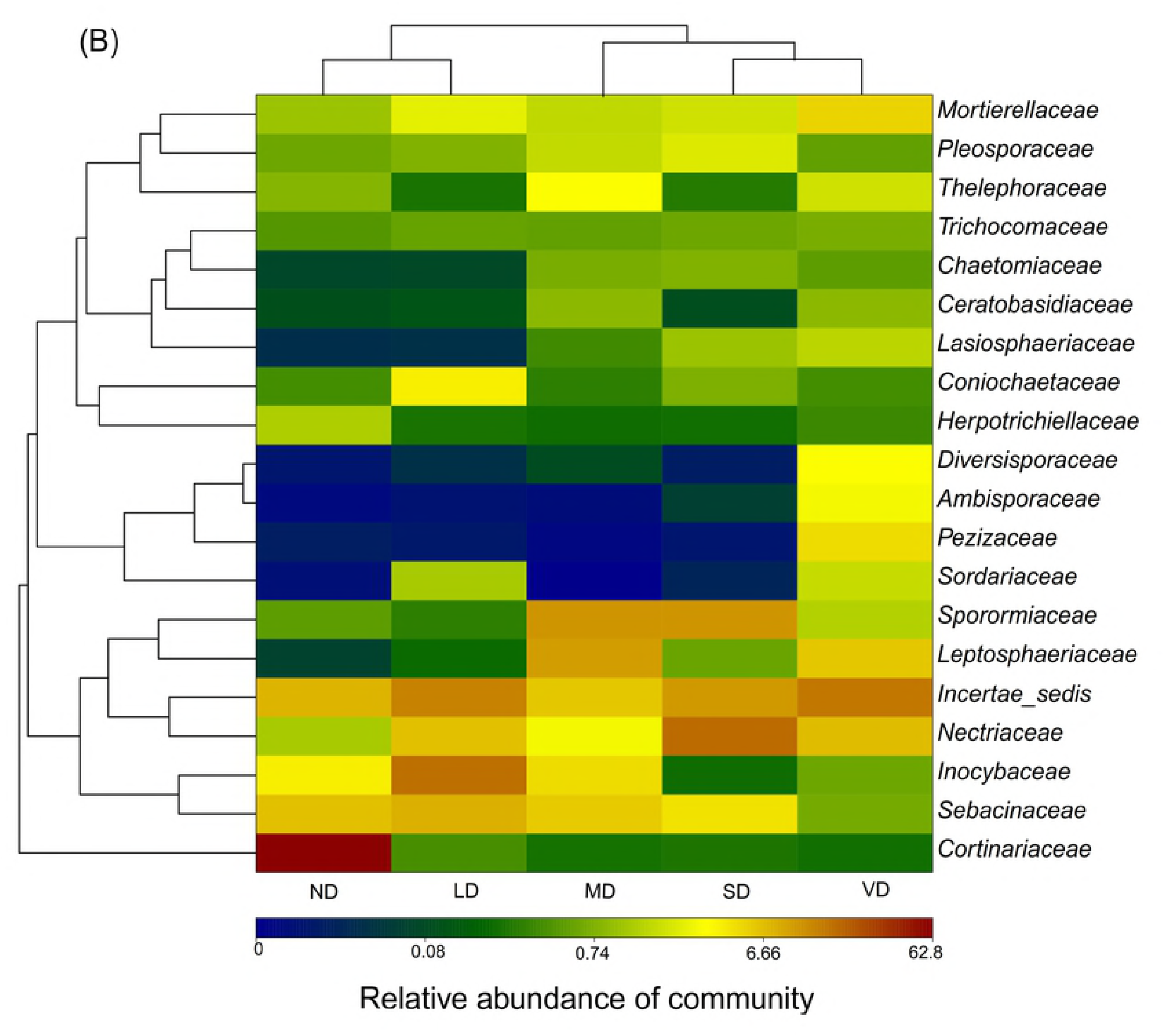
Heatmaps representation and cluster analysis of the microbial community among five samples.

**Fig 5.**
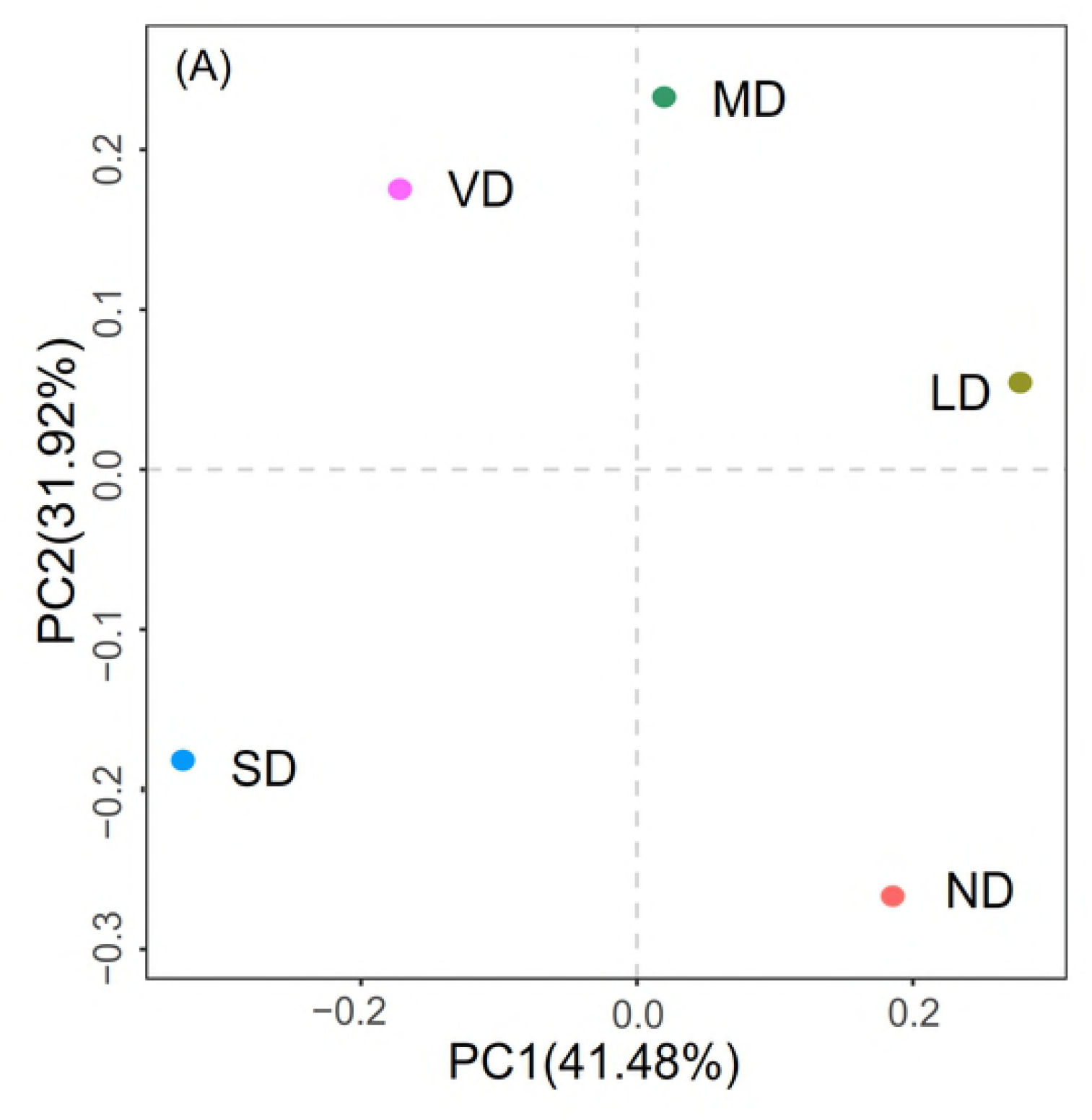
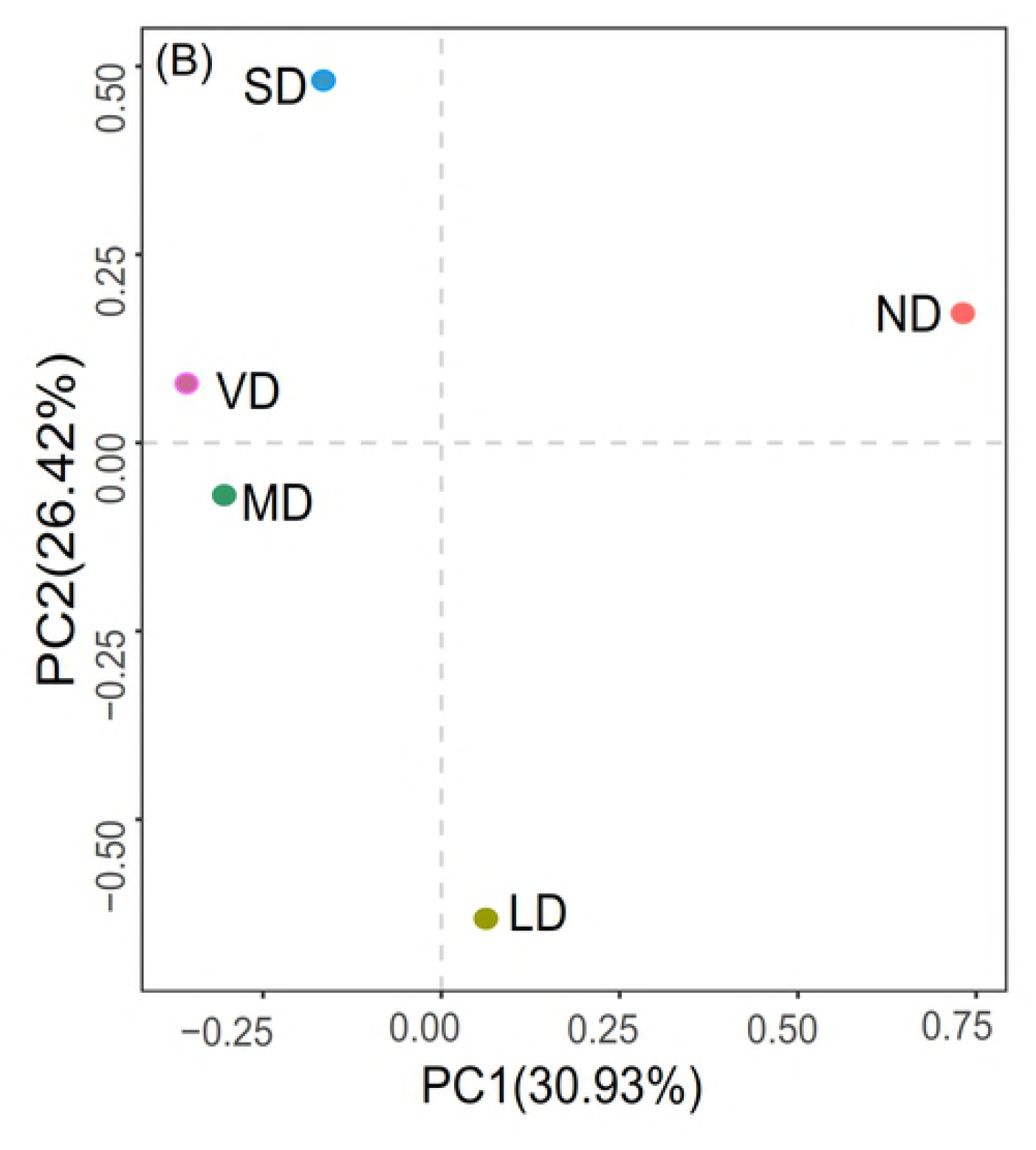
The results of bacterial (A) and fungal (B) communities for the five mixed samples according to the principal components analysis (PCA).

For fungi, the heatmap (Fig 4B) was based on the top 20 fungal families. The figure shows a two cluster sample division on a family rank level: ND and LD into one group, and SD, VD, and MD into another. The PCA score plot (Fig 5B) was in agreement with the heatmap, indicating high fungal community similarity between ND and MD and between MD, SD, and VD. ND and LD grouped to the right, and MD, SD, and VD grouped to the left of the graph along the PC1 axis, with a total variance of 30.93%. The PC2 axis accounted for 26.42% of the total variances.

### Correlation between community structure and environmental factors

The Monte Carlo test showed that the total explanatory powers of measure variables explained 44.5% and 35.2% of the total variation in bacterial communities on the first and second axis, respectively (Fig 6A). Results showed that soil bacteria under different degradation stages was significantly correlated to soil NO_3_^-^-N (*P* <0.01), SOC (*P* <0.05), TN (*P* <0.05), SM (*P* <0.05), and BGB (*P* <0.05), whereas soil NH_4_^+^-N, BD, AGB, etc. (*P* >0.05) had no obvious effect on bacteria (S2 Table).

**Fig 6.**
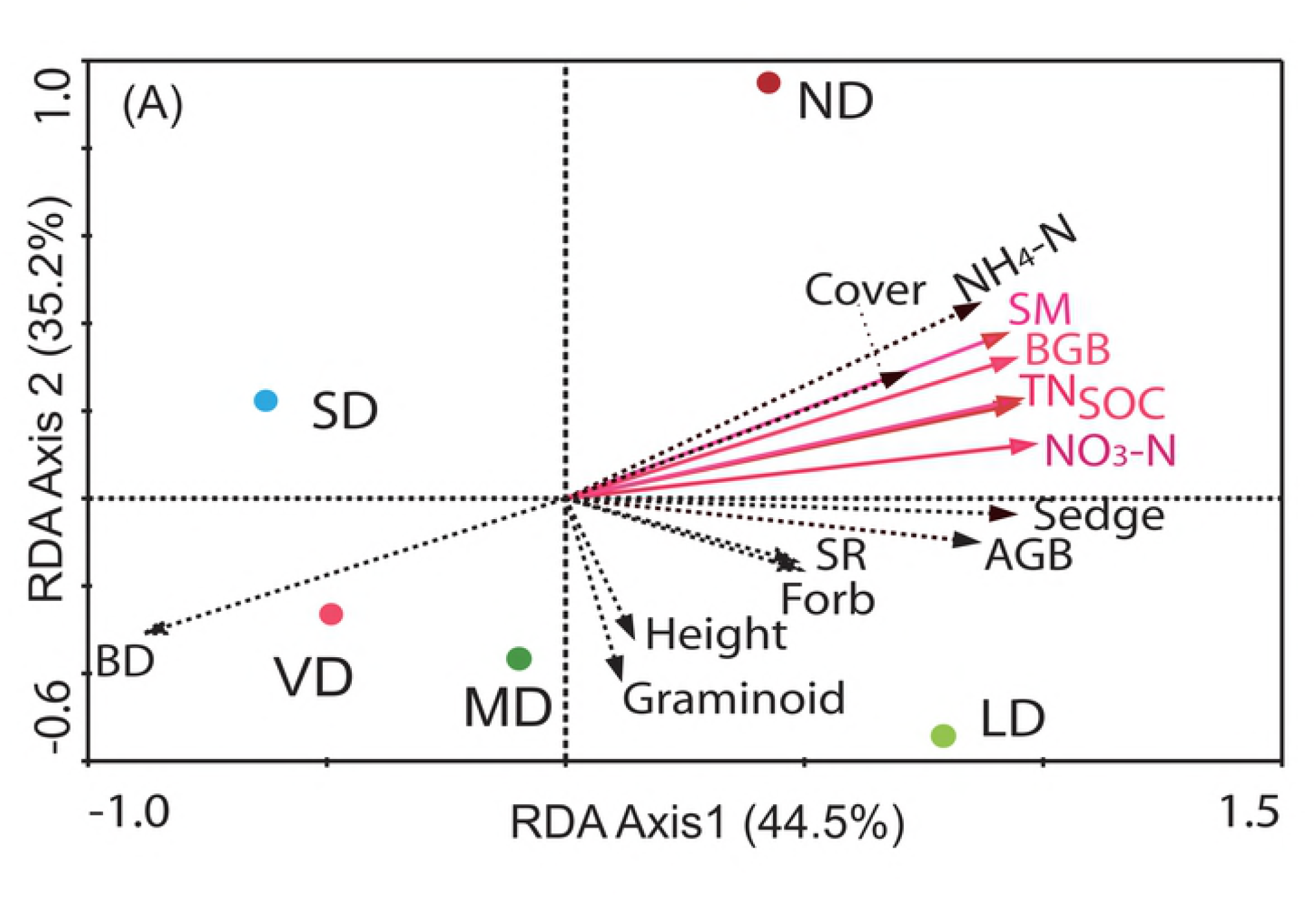
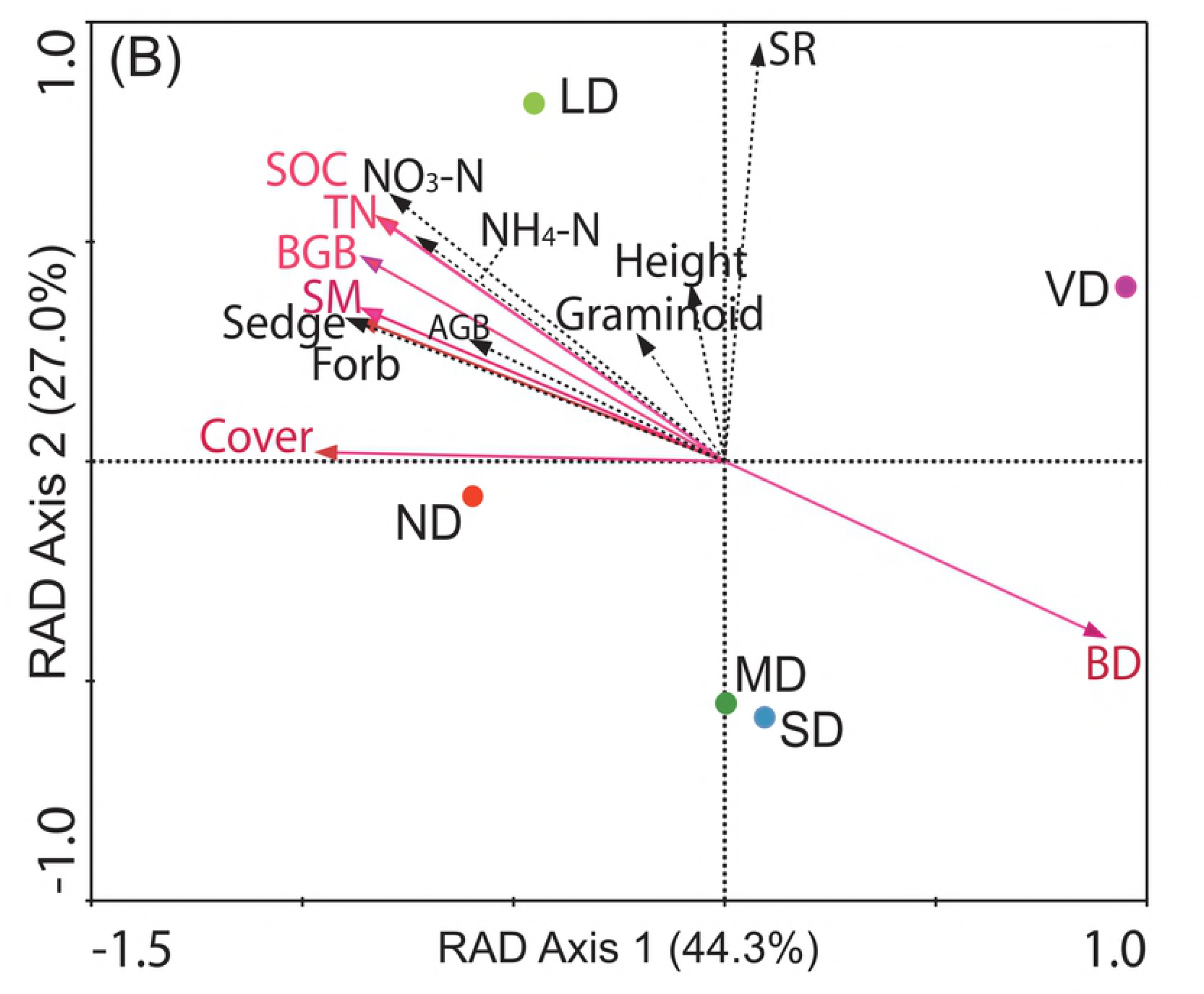
Relationships between soil and plant properties (arrows) and bacterial (A) and fungal (B) community structure according to the redundancy analysis (RDA). (A red solid arrow line means P<0.05; A black dashed line means P>0.05).

For fungi, the Monte Carlo test showed that the total explanatory powers of the measured variables explained 44.3% and 27.0% of total variation in fungal communities on the first and second axis, respectively (Fig 6B). Results showed that soil fungi were significantly correlated to soil TN (*P* <0.05), SOC (*P* <0.05), SM (*P* <0.05), BD (*P* <0.05), BGB (*P* <0.05), and coverage (*P* <0.05) whereas soil NO_3_^-^-N, NH_4_^+^-N, AGB, etc. (*P* >0.05) had no obvious effect on bacteria (S2 Table).

## Discussion

This study investigated how soil bacterial and fungal communities vary with changes in specific soil and plant properties in the QTP while discerning which factors significantly influence associative microbial community structure with alpine meadow degradation.

### Soil microbial community characteristics and its relationship with biotic and abiotic factors

Consistent with previous studies [19,21], increasing meadow degradation in our study significantly decreased the soil nutrient status (Table 3). This was due to a decrease in plant coverage and biomass, a reduction in SM, and an increase in soil BD in conjunction with degradation severity (Tables 1 and 3), which could potentially lead to nutrient leaching and contribute to nutrient loss [31]. In our study, both the bacteria and fungi genera heatmaps showed a two-cluster sample division. The first cluster was the ND and LD group, and the second cluster was the SD and VD group, which first clustered together before clustering with MD, resulting in the second SD, VD, and MD group. These results could be explained by a significant change in plant (i.e., coverage and BGB) and soil (i.e., SOC, TN, NO_3_^-^-N, NH_4_^−^-N, SM, and BD) factors in MD, SD, and VD compared to ND, while ND and LD exhibited no significant difference in the above factors (Tables 1 and 3).

About 79.7% and 71.3% of variation in bacterial and fungal composition could be explained by soil physicochemical properties and plant characteristics, respectively (Fig 6A, B), indicating that soil nutrient (i.e., SOC and TN) and moisture content, plant coverage, and BGB could potentially be key factors in defining changes in microbial composition under conditions of meadow degradation. Similar to our results, previous studies have demonstrated that the factors listed above affect soil microbial communities [32–34]. Variability in SM may reduce soil water availability averages [35], leading to reduced soil microbial community C use efficiency [36] and ultimately shifting the biomass or ratios of fungi and bacteria in soil [37]. At present, a number of studies have focused on the effect of nitrogen (N) gradients on soil microbial communities. This is because ecosystems around the world are being subjected to elevated levels of N [38,39]. For example, Ramirez et al. (2012) [40] found that N fertilization significantly affects total bacterial community composition. Conversely, increased N fertilizer dosages could potentially have a negative impact on C cycling in soil while, at the same time, promoting fungal genera with known pathogenic traits [38]. Moreover, the significant (*P* < 0.05) correlation found between bacterial and fungal communities and plant BGB (S2 Table) demonstrates that degradation-induced changes in plant species composition and biomass also exert strong effects on microbial communities. Wallenstein et al. (2007) [32] also suggested that plants strongly regulate microbial communities through the role they play in substrate supplies (e.g., litter, root turnover, and exudates) and by changing the physical environment in the active soil layer.

### Bacterial and fungal community response to meadow degradation

The bacterial phyla in meadow soil investigated in this study exhibited low variability in the different samples (Fig 2A); however, the composition of fungal phyla significantly changed under conditions of degradation severity (Fig 3A), indicating that fungal communities are more sensitive to degradation than bacterial communities. It was also reported that biotic and abiotic factors have a greater influence on fungi than bacteria [41]. The higher sensitivity of fungi that have been reported by some recent studies suggests that soil fungal communities are highly responsive to changes in SM and soil nutrient limitations than bacteria [42–44].

If fungal groups differ in their preference to substrate utilization processes [45], degradation-induced effects on biogeochemical properties will cause marked changes in specific species [46]. In our study, we consistently found that the dominate *Ascomycota* community structure significantly increased with increasing levels of meadow degradation severity (Fig 3A). Growth rates of *ascomycetes* were correlated to N availability, while their activity may dramatically accelerate C decomposition [47]. Additionally, the abundance of *Basidiomycota* significantly decreased in response to meadow degradation (Fig 3A). *Basidiomycetes* are widely recognized as lignin decomposers [48], and its capacity to utilize this recalcitrant substrate will likely hinder the development of this fungal group given that we detected a reduction in plant litter (biomass) with an increase in meadow degradation severity (Tables 1 and 2). Specifically, members of the *Cortinariaceae* family dominated the Basidiomycota phylum, and the proportion of reads exhibited high variability in abundance in the different samples; namely, in decreasing order of abundance, ND (62.80%), LD (0.42%), MD (0.23%), SD (0.26%), and VD (0.22%). Therefore, the association between fungi and plants may be responsible for the higher sensitivity of fungal composition to degradation severity compared to the bacterial composition.

For the bacterial groups, the overall proportion of *Proteobacteria* generally decreased while *Actinobacteria* generally increased in MD and VD compared to ND and LD (Fig 2A). This could have been because many members of *Proteobacteria* (particularly *α-proteobacteria*; Fig 2B) prefer nutrient-rich environments, whereas *Actinobacteria* are well adapted to oligotrophic environments [49,50]. Therefore, observed shifts are consistent with a decrease in the status of soil nutrients (i.e., C and N content) and the degradation of permafrost. Thus, degradation-induced environmental changes exert effects on microbial communities.

## Conclusions

In conclusion, this study, using the high throughput Illumina MiSeq sequencing method, provided a detailed description of variation in alpine meadow bacterial and fungal communities under different levels of degradation severity. For bacterial communities, these differences likely resulted from combined differences in soil properties and plant characteristics (most closely associated with SOC, TN, SM, NO_3_^-^-N, and BGB) rather than to single biotic or abiotic factors that combined to create the unique characteristics representative of each site. On the other hand, changes in soil fungal communities were mainly attributed to variations in SOC, TN, SM, BD, BGB, and plant coverage. Although this study will aid in our understanding of changes in soil microbial communities throughout the whole alpine meadow degradation process, further research into the mechanisms that underlie our findings would also be of great interest.

## Acknowledgements

The study was financially supported by the National Key Research and Development Program of China (2016YFC0501803) and the National Science Foundation of China (41771229, 41771233).

## Supporting information

**S1 Table**. The results of MiSeq sequencing and α-diversity estimates of the five mixed samples.

**S2 Table**. Correlations between environmental factors with the ordination axis and significance test of single variables for the bacteria and fungi community structure obtained from the RDA results.

